# Targeting caveolae to pump bispecific antibody to TGF-β into diseased lungs enables ultra-low dose therapeutic efficacy

**DOI:** 10.1101/2022.09.12.507679

**Authors:** Anil H. Kadam, Kathirvel Kandasamy, Tim Buss, Brittany Cederstrom, Chun Yang, Sreekanth Narayanapillai, Juan Rodriguez, Michael D. Levin, Jim Koziol, Bogdan Olenyuk, Zea Borok, Adrian Chrastina, Jan E. Schnitzer

## Abstract

The long-sought-after “magic bullet” in systemic therapy remains unrealized for disease targets existing inside most tissues, theoretically because vascular endothelium impedes passive tissue entry and full target engagement. We engineered the first “dual precision” bispecific antibody with one arm pair to precisely bind to lung endothelium and drive active delivery and the other to precisely block TGF-β effector function inside lung tissue. Targeting caveolae for transendothelial pumping proved essential for delivering most of the injected intravenous dose precisely into lungs within one hour and for enhancing therapeutic potency by >1000-fold in a rat pneumonitis model. Ultra-low doses (μg/kg) inhibited inflammatory cell infiltration, edema, lung tissue damage, disease biomarker expression and TGF-β signaling. The prodigious benefit of active vs passive transvascular delivery of a precision therapeutic unveils a new promising drug design, delivery and therapy paradigm ripe for expansion and clinical testing.

## Introduction

Modern medicine seeks precision in diagnosis and therapy. Ehrlich’s ideal of “aiming precisely” using drugs remains a challenge (1-6). Molecular medicine has advanced to achieve unparalleled interactive specificity in the high affinity binding of imaging and therapeutic agents. But this target binding precision does not automatically translate into the low dose therapeutic efficacy expected with high specific affinity. Many antibodies and other drugs designed to precisely target a therapeutic moiety expressed inside key diseased organs like lung actually do so below theoretical expectation after systemic administration (2-6). Targeting and molecular imaging are slow, inefficient and not particularly specific. Frequently it is not the target choice nor binding specificity or affinity but rather poor access that is limiting precision-targeted therapies *in vivo*.

A thin monolayer of endothelial cells (ECs) lining the inner surface of all blood vessels physically separates drugs circulating in the blood from tissue cells (3, 4). Systemic therapeutics must frequently overcome this major biological barrier to gain target access inside key organs and attain effective concentrations locally at the disease site. Currently all drugs depend on passive transvascular transport, primarily by diffusion and hydraulic drag through intercellular EC junctions or gaps, to reach and bind their targets inside tissue (4-6). Nonspecific and inefficient penetration of solid tissues necessitates high, sometimes near toxic doses to drive enough drug uptake to realize therapeutic efficacy. The current passive drug delivery paradigm appears to likely limit both therapeutic potency and efficacy while promoting undesired on- and off-target adverse effects elsewhere in the body (2-6).

Advances in genetic antibody engineering in the last few years have permitted designing a whole new generation of bi- and multi-specific antibodies to address the unmet medical need for better targeting and therapeutic precision without deleterious side effects. In theory dual binding can improve tissue targeting and therapeutic potency, efficacy and index of standard monospecific antibodies. But bispecifics still rely on passive delivery and thus have achieved their biggest gains so far in treating hematogenous disease where therapeutic targets are inherently more accessible. Lowering doses could benefit patients by minimizing toxicities usually seen at high doses and improving greatly the therapeutic index. In many cases, we don’t know whether drugs have achieved optimal concentration for maximum therapeutic effect, and we have to wonder, when drugs fail, whether they reached the target, especially inside solid tissue, at the necessary concentration to get full impact. Given that modern drugs are so well made and tested for maximal pharmacological activity, then with proper delivery assured the target must be lacking, for instance in expression homogeneity or in driving the pathology. Here we engineer a bispecific quad antibody format to create the first precision therapeutic to go beyond passive transvascular transport into a solid tissue, the lung.

Our group discovered an alternative and promising, active and specific, caveolae-mediated transcytotic mechanism in endothelium in vivo (7-10). We established that caveolae are not just static invaginations but can actually be dynamic transport vesicles with the molecular machinery for vesicular fission and fusion (11-15) necessary to carry and even pump select molecular cargo across microvascular endothelium. Proteomic mapping of the EC luminal surface and caveolae revealed apparent tissue-specific targets (8). Ultimately our imaging studies including intravital fluorescence microscopy of lung showed that select imaging probes, only when binding a target highly concentrated in lung EC caveolae, such as aminopeptidase P2 (APP2), can be transported rapidly across the EC barrier to concentrate solely in the lung interstitium within minutes of intravenous (iv) injection; this extravasation continued even against a concentration gradient which by definition constitutes active transport or pumping (7, 10).

Although clearly promising in theory, whether and to what degree targeting this caveolae-mediated transendothelial pumping system can help solve the long-standing drug delivery problem remains debatable. How much the EC barrier contributes on its own is unknown. Can an authentic precision therapeutic with remarkable high affinity and documented efficacy benefit from active delivery and in what way and how much? Here, we begin to answer these critical questions and discover and quantify a striking therapeutic impact.

We exploit the active precision delivery portal for possible therapeutic utility by genetically engineering a new antibody class, the first bispecific quad antibody where two arms enable active transvascular delivery into a normally restrictive solid tissue so that the second pair of arms can block the therapeutic target deep inside the solid tissue. We recombinantly engineer this “dual precision” therapeutic to combine key binding domains of a precision therapeutic antibody specifically neutralizing transforming growth factor-β (TGF-β) effector function with a precision lung-targeting and penetrating antibody specific for APP2. Testing the effects of active vs passive transvascular delivery on a precision therapeutic is now possible for the first time.

TGF-β regulates inflammatory and tissue remodeling responses that direct many acute and chronic pathologies in lung and other organs (16-20). Inhibiting TGF-β with small molecules and antibodies shows impressive potency in treating many diseases including lung diseases ranging from acute respiratory distress syndrome (ARDS) to chronic fibrosis (21-25) and quite recently COVID-19 (26-28). But effective treatment requires repeated high doses (mg/kg) that can elicit significant on- and off-target tissue side effects, including cardiovascular toxicity, bleeding, and death (29-32). TGF-β’s pleiotropic functions and widespread tissue expression enables further consequences from disrupted TGF-β signaling: pathogenic immune response, cancer, abnormal wound healing, neurodegenerative diseases, pulmonary hypertension and renal and cardiac fibrosis (33). Here we attempt to render lung targeting precision to a TGF-β blocking antibody for treating lung disease.

## Materials and methods

### Animals

Female Sprague Dawley rats (ENVIGO, USA) weighing 200-220 g were housed at 25°C with a 12 h day/night cycle and food and water *ad libitum*. Animals were maintained under these conditions until the end of the study. Care and experiments were approved by the PRISM Institutional Animal Care and Use Committee.

### Antibodies and reagents

Recombinant rat aminopeptidase 2 (APP2) as well as recombinant antibodies fresolimumab, 833c, 833c&Freso and 833cX&Freso were genetically engineered in-house.

### 833c&Freso genetic engineering and characterization

833c is a chimeric antibody with murine variable regions (VH and Vk) and human constant regions (CH1-CH2-CH3 and Ck) (*9*). The heavy chain of 833c&Freso was synthesized by attaching the single chain Fv fragment of human fresolimumab to the C-terminus of the human CH3 domain via a SGGGGS linker. The whole cassette was cloned into the TOPO-TA cloning vector pcDNA3.4. The light chain was encoded by a separate pcDNA3.4 vector. Antibodies were transiently expressed in CHO-S cells using the Expifectamine transfection kit at a molar ratio of heavy to light chain plasmids of 1:1.2. ExpiCHO enhancer and feed were added on day 2. Cells were typically harvested on day 8 or 9 post transfection and removed from the culture by centrifugation. The supernatant was filtered through a 0.2 μm sterile filter unit and antibody purified with a protein G Sepharose column (Cytiva HiTrap) on an AKTA purifier. Batch to batch stability was controlled using capillary isoelectric focusing.

### Commercially available test kits and reagents

Rabbit anti-Phospho-Smad2 (Ser465/467)/Smad3 (Ser423/425) (D27F4), rabbit anti-Galectin-3/LGALS3, rabbit anti-Phospho-Akt (Thr308) (D25E6), rabbit-anti-Akt (pan) (C67E7), rabbit Phospho-p70 S6 Kinase (Thr389) (108D2), rabbit anti-p70 S6 Kinase (49D7), rabbit anti-PDGF Receptor β (28E1), and rabbit anti-GAPDH (D16H11) XP were purchased from Cell Signaling Technology (Danvers, MA). We used rabbit polyclonal anti-Fibronectin (ab2413) (Abcam, Cambridge, MA), recombinant hu TGF-β1(ProSci, Poway, CA), anti-TGF-β1 IgY-Biotin (R&D Systems, Minneapolis, MN), anti-hIgG-HRP (Jackson ImmunoResearch Inc., West Grove, PA), 1D11 (Invitrogen) and goat anti-rabbit IgG-HRP (Thermo Fisher Scientific, Carlsbad, CA). Rat Quantikine ELISA kit for TGFβ1, IL6, IL1β, WISP-1, CINC-1 and TIMP-1 were purchased from R&D Systems (Minneapolis, MN).

The following reagents were used: Bleomycin, USP grade (Teva pharmaceuticals, North Wales, PA or Hospira pharmaceuticals, Lake Forest, IL), CHAPS hydrate and protease inhibitor cocktail (Sigma-Aldrich, St. Louis, MO), and phosphatase inhibitor cocktail II (Alfa Aesar, Tewksbury, MA). Nitrocellulose membrane filter paper sandwich (Invitrogen), Novex 8–16% Tris-Glycine-gel system (Invitrogen), SuperBlock™ (PBS) blocking buffer, 96-well flat-bottom microtiter plates, bovine serum albumin, 10X PBS pH=7.4 (Gibco), HPR-conjugated streptavidin (Pierce), SuperSignal™ West Pico PLUS chemiluminescent substrate were purchased from Thermo Fisher Scientific (Carlsbad, CA). ABTS substrate (KPL, Maryland WA), TRIS-buffered saline (TBS, 10X) pH=7.4 for western blot (Alfa Aesar, Tewksbury, MA) and 10% neutral buffered formalin (Leica biosystems, Buffalo Grove, IL) were used. Gel and western blot images were captured using a Konica Minolta Bizhub c284e scanner.

### Rat APP2 binding assay

We coated 96-well flat-bottom plates with recombinant rat APP2 (5 μg/ml) overnight at room temperature. After washing 3 times with PBS containing 0.05% Tween-20 (PBST), plates were blocked with PBS containing 1% BSA and 0.01% Tween-20 for 90 min. Serially diluted antibodies were added, and plates were incubated for 2 h, washed and incubated with anti-hIgG-HRP for 1 h. After final washing, ABTS substrate (100 μl/well) was added for 15 min, and the reaction was terminated with 0.1 M citric acid, pH=4.0 (100 μl/well). Absorbance was read at 415 nm and results analyzed using GraphPad software.

### TGF-β1 binding assay

We coated 96-well plates with titrated antibody solutions (210.0-0.002 nM) overnight at 4°C. After washing and blocking, 100 μl recombinant hu TGF-β1 (50 ng/ml) was added and incubated at 37°C for 2 h. Plates were washed and incubated with 100 μl anti-TGF-β1 IgY-Biotin (1.0 μg/ml) for 1 h. After washing, plates were incubated with HRP-streptavidin (1:2500). After final washes, ABTS substrate (100 μl/well) was added for 15 min, the reaction terminated with 0.1 M citric acid, pH=4.0 (100 μl/well) and absorbance read at 415 nm. Results were analyzed using GraphPad software.

### Bleomycin-induced acute lung injury

Bleomycin was dissolved in 1X PBS pH=7.4. On day 0, rats were anesthetized with inhaled isoflurane (3%–5%) and received a single instillation it of bleomycin (2 U/kg) using a 18 gauge needle attached to a 1 mL tuberculin syringe. Control animals received 300 μl 1X PBS. After instillation, rats were allowed to recover from anesthesia and returned to their cages with free access to food and water. Animals were sacrificed on day 3, and acute lung injury relevant endpoints were analyzed.

### Treatments with mono and bispecific antibodies

All antibodies were dissolved in 1X PBS pH=7.4. 833c&Freso (1–100 μg/kg), fresolimumab (10–3000 μg/kg), 833c (10 μg/kg) and 833cX&Freso (10 μg/kg) were administered as a single dose iv through the tail either prophylactically one h prior to bleomycin challenge (−1 h), or on day 1 (D1) or day 2 (D2), or day 1 and day 2 (D1&2). Control animals received PBS. Animals were sacrificed on day 3 (D3).

### SPECT/CT tomographic imaging

833c&Freso, 833cX&Freso, and fresolimumab were radiolabeled with iodine-125 (^125^I) (PerkinElmer, Waltham, MA) using Pierce™ Iodination Beads (Thermo Fisher Scientific, Carlsbad, CA) according to the manufacturer’s instructions, resulting in specific activities ranging from 5.5μCi/μg to 13μCi/μg. Radiolabeling did not affect apparent Kd values as determined by ELISA. SPECT/CT imaging was performed using a nanoScan SPECT/CT imaging system (Mediso, USA). Rats were injected iv with the indicated amount of antibodies 24 h after bleomycin treatment, and SPECT/CT images were captured after 1 h and 24 h. Note that the 1 h timepoint is an average, because the imaging process takes about 30 to 90 min. Images were acquired for 60 sec/projection using a four head gamma camera with a rat-seized multi-pinhole. Images were collected in a 360° orbit with 60 s sampling every 6°. The pulse height analyzer window was >28.40 keV with a width of 20%. SPECT/CT reconstruction was performed using Nucline nanoScan software (Mediso, Arlington, VA). CT voltage was set at 50 Kv, exposure for 300 msec and 720 helical projections were captured. After CTSPECT fusion, SPECT/CT 3-D data sets were processed with VivoQuant software (Invicro, Boston, MA). SPECT Planar imaging with ^125^I-833c&Freso was done at 3 μg/animal.

### Size-exclusion chromatography with multi angle light scattering detection (SEC-MALS)

Antibodies were concentrated to 2 mg/mL using Cytiva Vivaspin® 500 50kDa MWCO (Cytiva, United Kingdom) spin filters. A total volume of 50 μL was injected into an HPLC instrument (Nexera SIL-20AC XR autosampler (Shimadzu), Nexera LC-20AD XR pump (Shimadzu), WTC-030S5 size exclusion chromatography column (5 μM, 300 Å, 7.8 × 300 mm, Wyatt Technology, USA) stabilized at 25°C in a CTO-20AC column oven (Shimadzu), SPD-M20A diode array detector (Shimadzu) followed by MiniDawn TREOS light scattering and Optilab T-rEX refractive index detectors (Wyatt Technology, USA). The DLS signal was collected by the TREOS detector and processed by the DynaPro NanoStar detector (Wyatt Technology, USA). Separation was carried out under an isocratic flow at 0.5 ml/min of 150mM sodium phosphate buffer, pH 7.0. Data was processed using ASTRA® V 7.3 software (Wyatt Technology, USA).

### In vivo biodistribution analysis

One or 2 days after bleomycin administration rats were injected iv with 3 or 10 μg radiolabeled antibodies. After 1 or 24 h, rats were euthanized. Blood and excised major organs were weighed, and radioactivity was measured on a Wizard 1470 Automatic Gamma counter (PerkinElmer Life Sciences, Wallac Oy, Finland). Radiation uptake is expressed as percentage of injected dose (% ID) and percentage of injected dose per gram of tissue (%ID/g). Tissue targeting index (TTI) was calculated as antibody per g of lung tissue/antibody per g of blood. We observed no significant differences in results from day one or two and 3 or 10 μg, and data were pooled.

### Bronchoalveolar lavage (BAL)

Three days after bleomycin instillation, animals were euthanized with Euthasol (100–120 mg/kg) by intraperitoneal injection. The trachea was exposed following a small incision to the skin and BAL was performed 3 times using a plastic cannula with 2 mL 1X PBS (pH=7.4). Volumes of individual BAL aspirates were pooled.

### Assessment of pulmonary inflammatory cells

We mixed equal volumes of BAL fluid (BALf) and Turk’s solution and counted total leukocytes manually using a hemacytometer (Hausser Scientific, Horsham, PA). The remaining fluid was centrifuged at 4000 RPM for 5 min at 4°C, aliquots of BALf supernatant were collected aseptically and stored at -80°C until analysis. Cell pellets were reconstituted in rat serum and stained with Leishman solution on frosted glass slides (Leica Biosystems). Using a light microscope (BX2, Olympus, Tokyo, Japan) at 100X magnification 500 cells/slide were counted. Cells were categorized based upon morphology into neutrophils, lymphocytes, eosinophils, or macrophages.

### Lung harvest for protein-, histology- and edema analysis

After BAL, right lungs were harvested from animals, washed in 1X PBS and stored at -80°C until western blot analysis. For histology, left lungs were carefully removed and stored in 10% neutral buffered formalin (NBF). Paraffin-embedded tissue (4 μm slides) was stained with haematoxylin and eosin. In a separate set of rats, lungs were harvested without performing BAL to determine the severity of pulmonary edema by the ratio of whole wet to dry weight. To obtain dry weight, lungs were incubated in an oven (45°C for 24 h).

### Lung injury scoring criteria

Pathological changes in lung tissue were assessed using the following criteria which were adapted from previously published protocols (34-36): 1. cell infiltration severity 0-4 (0: normal, 1: mild, 2: moderate, 3: severe, 4: highly severe), 2. presence and absence of hyaline membranes (0: absence and 1: presence), 3. presence and absence of pulmonary edema (0: absence and 1: presence), 4. alveolar septal thickening 0-3 (0: Normal,1: 2X, 2: 3X, 3: >3x) in total 20 random microscopic fields (20X magnification). Lung injury scores reflect the sum of criteria above. Lung samples for histology were obtained from 3 different independent experiments (n=2-3/group).

### Quantification of biomarkers of inflammation in rat lungs

TGF-β, IL-6, IL-1β, WISP-1, CINC-1, and TIMP-1 levels in BALf were determined using rat Quantikine ELISA following manufacturer’s instructions.

### Lung preparation and western blot analysis

Lungs were placed in 1 ml PBS containing 0.1% (v/v) protease and phosphatase inhibitor cocktail and stored at -80°C until use. Lungs were homogenized on ice in 5 mL CHAPS buffer containing protease and phosphatase inhibitors using a homogenizer (PT3100-POLYTRON, Kinematica, NY). Homogenates were centrifuged at 15,000 rpm for 15 min at 4°C, and the supernatant was stored at -80°C until use. Total protein concentrations were determined using bicinchoninic acid (BCA) assay. Proteins (50 μg) were separated on Novex 4–12% Tris-Glycine gels, transferred onto nitrocellulose membranes, blocked with SuperBlock™ for 2 h at room temperature (RT) and incubated with primary antibodies diluted in TRIS buffered saline pH=7.4 (TBS) containing 0.1% Tween-20 following manufacturer’s instructions. After washing with TBS-Tween, membranes were incubated with goat anti-rabbit IgG HRP-linked secondary antibody (1:10,000) for 1 h at RT and proteins were visualized by enhanced chemiluminescence.

### Statistical Analysis

All data are presented as mean ± S.D. Student’s t test was performed for comparing 2 groups and one way ANOVA followed by Dunnett’s multiple comparison test for more than 2 groups. Differences were considered statistically significant for P < 0.05 compared with bleomycin control.

### Data and materials availability

All data are available in the main text or the supplementary materials.

## Results

To ascertain the extent to which the EC barrier limits therapeutic impact and to overcome high dosing and poor lung-specific delivery, we generated a bispecific antibody by genetically splicing a single chain Fv (scFv) fragment of the TGF-β-neutralizing antibody fresolimumab onto the C-terminus of a lung-targeting APP2 chimeric antibody (833c) (Figs 1A and B). The two arms of the antibody, 833c and fresolimumab, were chosen to serve two distinct purposes; 833c to bind to EC caveolae and drive active delivery into the lung interstitium and fresolimumab to block TGF-β signaling pathways. This bispecific antibody 833c&Freso was expressed and purified up to 99% monomer (Figs 1C and D). Every batch of antibody was confirmed to be at least 97% pure. ELISA confirmed 833c&Freso maintained binding affinity to both targets (Figs 1E and F) unlike a mutated version 833cX&Freso, which did not bind APP2 by design (see Methods). Size-exclusion chromatography with multi angle light scattering detection confirmed only a minimal increase in the average hydrodynamic radius of the bispecific antibody 833c&Freso (6.1 nm) and 833cX&Freso (6.1 nm) compared to 833c (5.2 nm) and fresolimumab (5.0 nM) (Fig S1). Through recombinant engineering, we hope to have designed the first bispecific therapeutic to be pumped across the EC barrier to concentrate in the interstitium and engage its target deep inside a single solid tissue, in this case the lung. The delivery arms of the bispecific should by design attain the critical access for the therapeutic arms to reach and engage their target for utmost efficacy.

**Figure. 1.**
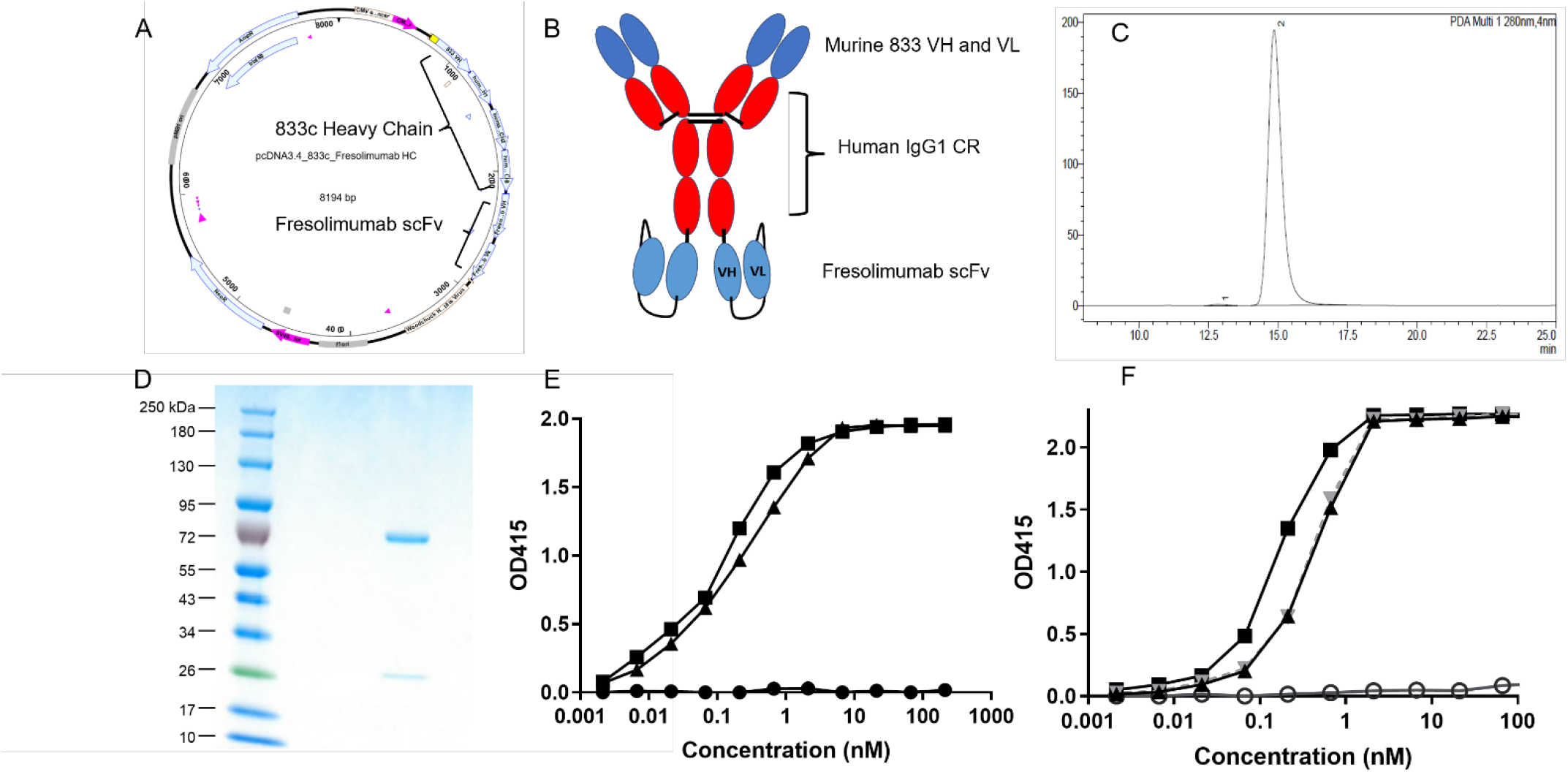
Genetic engineering and characterization of 833c&Freso. (**A**) Plasmid map for 833c&Freso heavy chain expression. Fresolimumab single-chain variable fragment (scFv) was cloned into the 3’ end of the 833c CH3 domain. (**B**) 833c&Freso schematic: scFv is attached to the CH3 domain at the 3’ end. Red: human domains, blue: murine domains. CR: constant region, VH and VL: variable regions. (**C**) Size exclusion chromatography and (**D**) Reducing SDS-PAGE of purified 833c&Freso. (**E**) ELISA binding to rat aminopeptidase P2 of 833c&Freso (triangles), 833cX&Freso (circles) and 833c (squares) (see Methods). (**F**) ELISA binding to TGF-β1 of 833c&Freso (triangle, black), 833cX&Freso (triangle, gray), fresolimumab (squares) and huIgG4, a isotype-matched control IgG for fresolimumab (open circles) (see Methods).

To test the therapeutic potential of this novel heterobifunctional antibody, we chose the clinically relevant bleomycin model. Bleomycin is used to treat cancer patients but can cause acute lung injury, fibrosis, ARDS and even death. In animal models, bleomycin is widely used to study lung inflammation and fibrosis (36). The bleomycin lung injury model shares mechanisms, pathology and sequelae seen in COVID-19 patients (25, 37, 38).

First, we tested lung immunotargeting of our antibodies. We challenged rats intratracheally (it) with bleomycin (see Methods) and after 24 or 48 h injected ^125^I-833c&Freso intravenously (iv). Whole body imaging showed similar robust lung uptake. Planar γ-scintigraphy and SPECT/CT in maximum intensity projection (MIP) as well as cross sections all revealed remarkably precise lung imaging, both 1 h and 24 h post injection (Fig 2A-B). Neither the mutant ^125^I-833cX&Freso nor ^125^I-fresolimumab, which both lack APP binding, targeted the lungs even after 24 h. They remained mostly in the blood and exhibited the expected non-specific blood distribution profile (Fig 2C-F).

**Figure 2.**
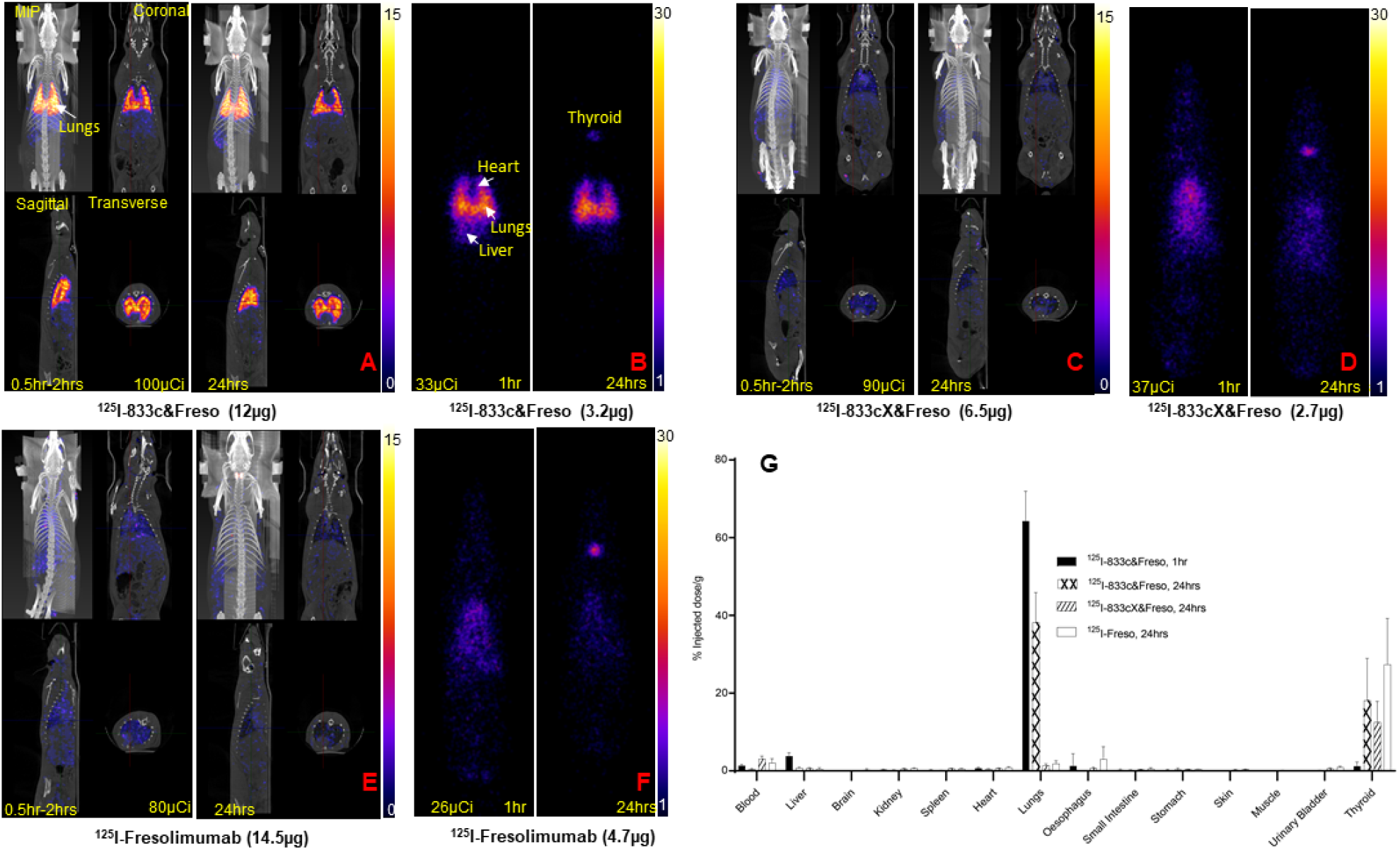
Lung targeting of 833c&Freso but not fresolimumab. After bleomycin treatment, rats were injected iv with radio-iodinated antibodies ^125^I-833c&Freso, ^125^I-833cX&Freso, and ^125^I-fresolimumab (see Methods). **(A, C, E)** SPECT/CT images of rats taken at 1* and 24 h. Maximum intensity projection (MIP), coronal, sagittal, and transverse cross sections are shown. **(B, D, F)** Planar γ-scintigraphy imaging at the indicated timepoints. A-F: N=2-3 rats per group **(G)** Biodistribution analysis of radioiodinated antibodies. Radioactivity was counted in the indicated organs excised from bleomycin-exposed rats. Uptake was quantified as the percentage of injected dose per gram tissue (means ± SD). N=3-10 rats per group. *Imaging started at 30 min and takes approximately 30 to 90 min.

Biodistribution analysis of excised organs confirmed lung immunotargeting with ^125^I-833c&Freso (64% of injected dose/g lung) and very low blood levels, all within 1 h (Fig 2G). Up to 83% of the total dose accumulated in the lungs. Blood clearance was very rapid in contrast to the other antibodies which just remained in the bloodstream, even after 24 h. As expected, dehalogenation of antibody facilitated obvious release of radioiodine with concomitant subsequent robust thyroid uptakes. Ultimately the two tissues with the most intense radioactive signal have a pumping mechanism: the lung via antibody transcytosis after APP2 binding in caveolae of lung ECs (7-10) and the thyroid through transmembrane pumping of freed radioiodine by the sodium-iodine symporter expressed in thyroid cells (39).

We quantified lung targeting precision by standard calculation of the tissue-targeting index (TTI) (antibody per g of lung tissue/antibody per g of blood) (10). 833c&Freso produced an impressive TTI of 117±31 (mean ± SD) whereas the TTI was less than 1 for the other antibodies, consistent with their poor lung targeting. TTI comparison proved that adding APP2 binding to fresolimumab in this bispecific format did indeed improve lung targeting precision by over 100-fold.

We used these targeting data to inform dose selection for therapy. Fresolimumab binds to all three TGF β isoforms, but TGF-β1 is predominantly induced during bleomycin lung pathogenesis (40). Knowing the apparent affinity constant for antibody binding to APP2 (∼0.2 nM) and fresolimumab to TGF-β1 (1 nM) (31, 41, 42), we must attain a few fold higher concentration first in the blood to enable robust pumping by caveolae and then inside the lung tissue to achieve enough TGF-β blockade to render therapeutic efficacy. Our imaging injection of 3 μg translates to ∼15 μg/kg dose; this generates peak blood levels above 1 nM, plenty for robust APP2 engagement and transport. With 70-80% delivered to the lung, we can attain ∼10 nM inside the lungs which is more than enough to drive TGF-β engagement and blockade. Using half log progression, doses of 1, 3, 10, and 30 μg/kg appeared warranted.

To assess pharmacological activity at μg/kg doses in vivo, we first examined prophylactic efficacy by injecting 833c&Freso iv 1 h before bleomycin challenge. The effects of dose escalation were evaluated on the inflammatory cell profile in bronchoalveolar lavage fluid (BALf) harvested on day 3. Bleomycin greatly increased total leukocyte, neutrophil and lymphocyte levels from normally low to nil levels (Fig S2). Rats treated with 833c&Freso doses above 3 μg/kg showed statistically significant reductions in total leukocytes and neutrophils (Fig S2).

We did not optimize these studies further but rather proceeded immediately to therapeutic testing. We selected day 2 post-bleomycin as most clinically relevant because bleomycin induced peak TGF-β1 expression near day 2 (Fig S3), the same day respiratory symptoms become clearly apparent. Again, total leukocyte and neutrophil levels were significantly reduced in animals receiving 1–30 μg/kg 833c&Freso on day 2 (Fig 3A and B). 10 μg/kg was statistically more effective than 1 μg/kg but statistically indistinguishable from 3 and 30 μg/kg. Inhibition of the bleomycin effect at 10 μg/kg averaged 54% for leukocytes and 57% for neutrophils (Fig 3B). Lymphocytes and macrophages appeared mostly unaffected.

**Figure 3.**
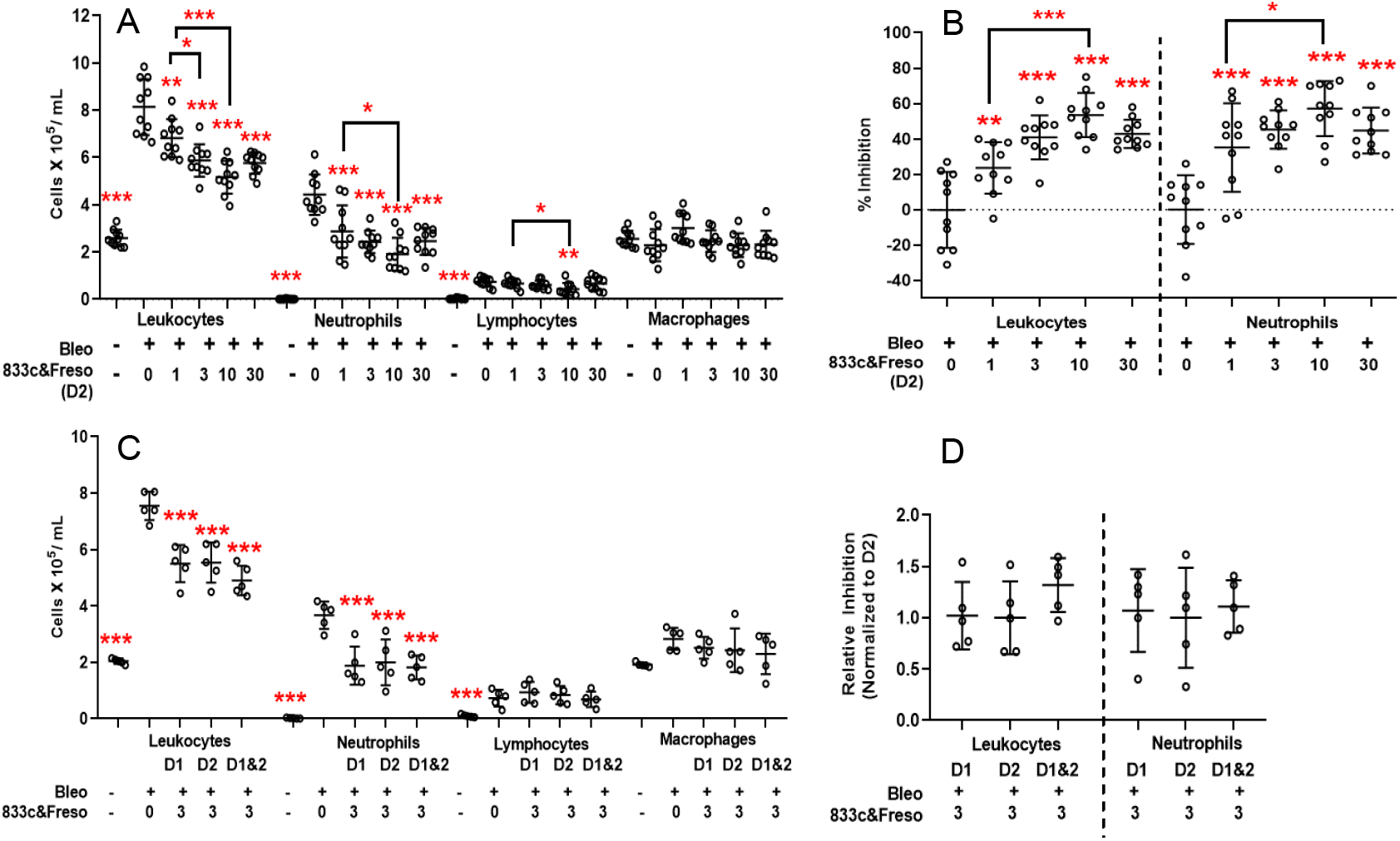
Therapeutic effects of 833&Freso on bleomycin-induced lung infiltration of inflammatory cells detected in BALf. 833c&Freso at the indicated doses (μg/kg) was injected iv into rats on the indicated days after bleomycin challenge **(A-D)** day 2 (D2), **(C-D)** day 1 (D1), **(C-D)** day 1 and 2 (D1&2)). BALf was collected on day 3. Total leukocyte and neutrophil count per ml of BALf **(A**,**C)** and percentage inhibition of the bleomycin effect on total leucocyte and neutrophil infiltration **(B)**. Data pooled from 2 independent experiments totaling n=10 rats per group. **(D)** Relative inhibition is normalized to single treatment on D2, with n=5 rats per group. Scatter graphs showing each data point with indicated means ± SD. *P< 0.03, **P< 0.01, and *** P< 0.001 vs. bleo only, by using one-way ANOVA followed by Dunnett’s test, except as indicated otherwise (A and B) when comparing dose differences with Student’s unpaired t test.

Next, we optimized dose regimen. We tested two injections at 3, 10 and 30 μg/kg given a day apart on day 1 and day 2. No obvious increase in efficacy was observed (Fig S4). A direct comparison in the same experimental cohort confirmed that there was no statistical difference with 3 μg/kg given on day 1 only, day 2 only, or day 1 and 2 (Fig 3C-D).

Histopathology confirmed that 10 μg/kg 833c&Freso administered on day 2 protects the lung tissue from bleomycin-induced inflammatory cell influx. After three days bleomycin caused obvious pneumonitis with rampant inflammatory cell infiltration in the tissue interstitium and alveolar air spaces as well as protein exudates and alveolar septal thickening (compare Fig 4A and B to 4E normal). These pneumonitis hallmarks (43) were ameliorated by treatment (Fig 4C and D). Lung injury scoring quantified the robust bleomycin effect (0.1±0.2 control vs 3.2±0.3 bleomycin) and its 41% inhibition by TGF-β blockade (1.9±0.2 bleomycin plus 833&Freso) (Fig 4F). In agreement, other experiments independently showed 833c&Freso reduced lung edema from bleomycin by 36%, as assessed by lung wet to dry ratios (Fig 4G).

**Figure 4.**
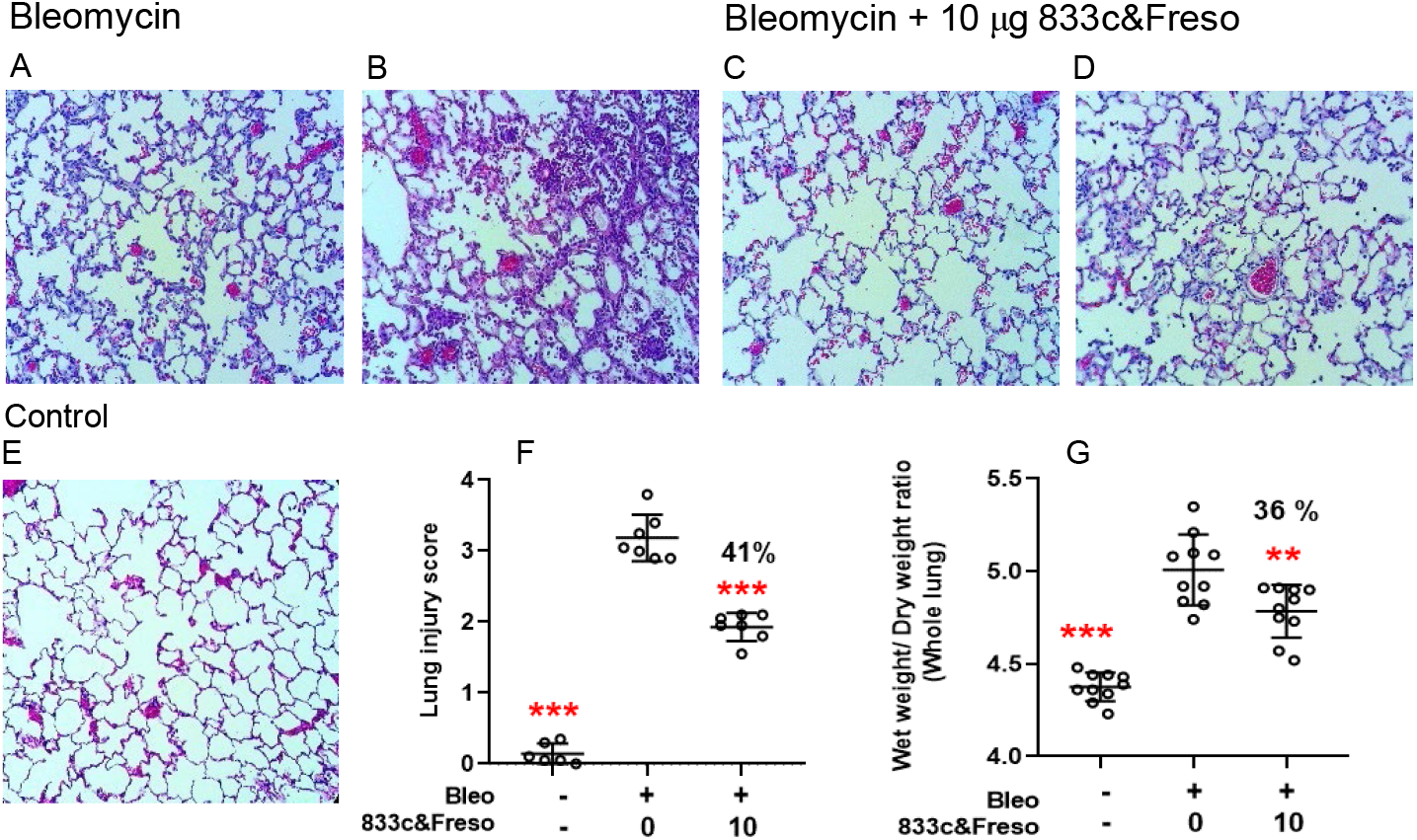
Therapeutic effect of 833c&Freso on lung pathology and edema from bleomycin. Rats were treated with 10 μg/kg 833c&Freso iv on day 2 after bleomycin challenge. Lungs excised on day 3 were sectioned and stained with H&E. Light microscopy images (×20) showing lung histopathology **(A-D)** vs. normal **(E)**. Tissue damage was clearly evident but varied within each group, ranging from low **(A, C)** to high **(B, D). (F)** Lung damage in each 20x microscopy field was scored as described in Methods. **(G)** Wet to dry lung weight ratio, n=10 total rats per group. Scatter graphs indicate means ± SD. Percentage (%) inhibition of lung injury scores **(F)** and edema **(G)** is indicated. **P< 0.01, and *** P< 0.001 were obtained vs bleo control by one-way ANOVA followed by Dunnett’s test.

Our literature search identified 12 well-known biomarkers of related lung diseases (18, 44-48), also induced by bleomycin (49), that we could readily and directly measure as proteins in BALf or whole lung homogenates from rats, using commercially available reagents. Although TGF-β1 levels appeared unaffected, 10 μg/kg 833c&Freso significantly decreased the bleomycin-induced signal in BALf for key interleukins, IL-6 (39%) and IL-1β (54%) (Fig 5 A); both are inflammatory cytokines regulated by TGF-β (50, 51) and induced by bleomycin (52). WISP-1, a well-known TGF-β downstream target (53) implicated in fibrogenesis (54), as well as the neutrophil chemoattractant CINC-1 and TIMP-1, a metalloproteinase inhibitor implicated in acute lung injury and tissue remodeling (55), were each reduced by 38%, 68% and 36%, respectively (Fig 5A).

**Figure 5.**
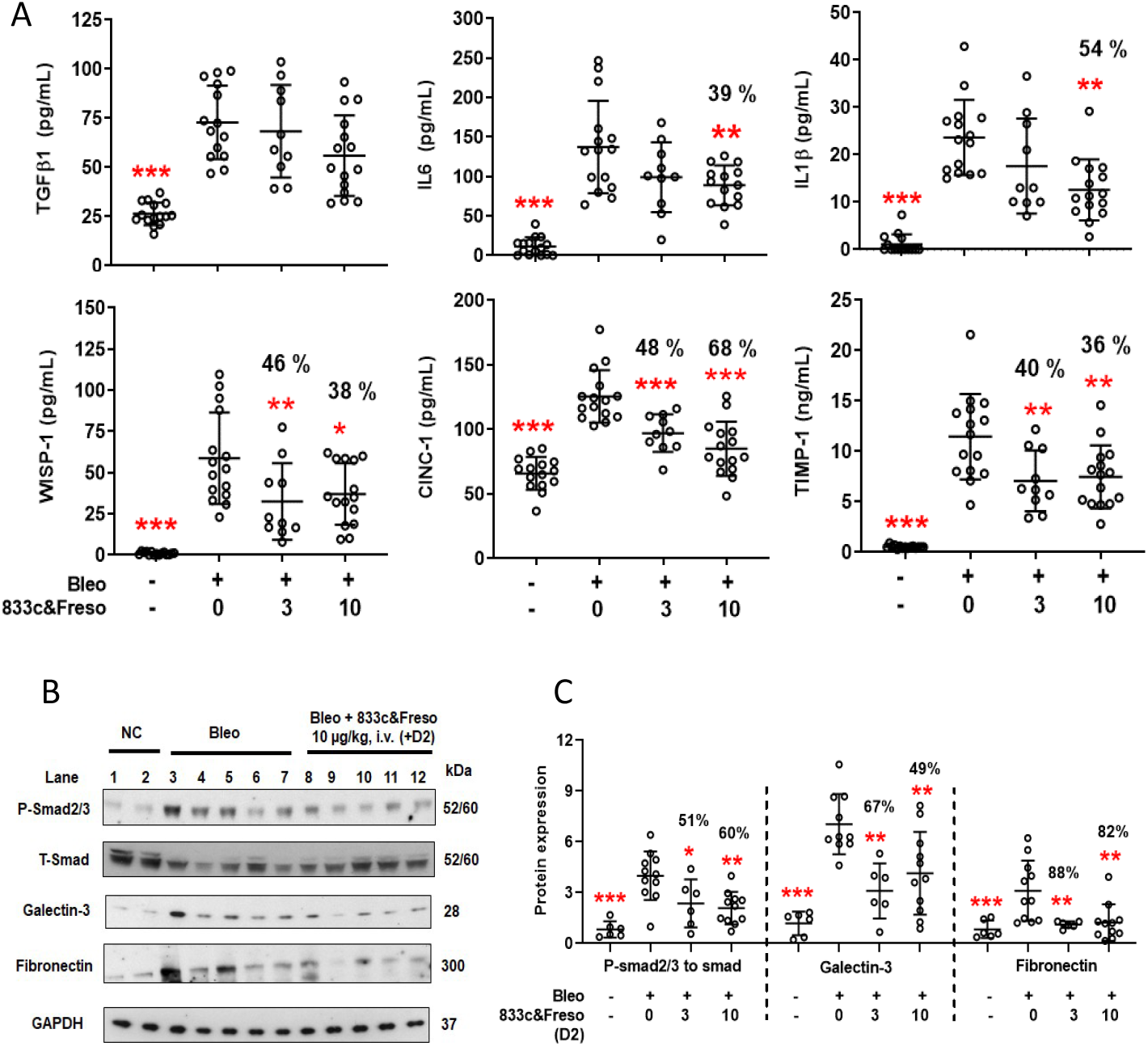
833c&Freso reduces TGF-β-mediated disease biomarkers and signaling in the lung. Biochemical assays were performed on lung samples taken from untreated rats and rats treated as indicated with 833c&Freso on day 2 after bleomycin challenge. BALf or lung tissue was harvested on day 3. **(A**) Concentration of the indicated biomarkers detected in BALf. Data pooled from 2-3 independent experiments with n=10-15 total rats per group. **(B)** Western blot of lung tissue homogenates using antibodies to the indicated proteins. GAPDH: loading control. Data pooled from 2-3 independent experiments with n=6-11 total rats per group. **(C)** Relative expression of indicated proteins derived from western blot image analysis. Smad 2/3 protein phosphorylation: ratio of P-to T Smad 2/3. Graphs show means ± SD. Percentage (%) inhibition of biomarkers in BALf **(A)** and proteins expressed in the lung **(C)** are shown. *P< 0.03, **P< 0.01, and *** P< 0.001 were obtained vs bleo control by one-way ANOVA followed by Dunnett’s test.

Western blot analysis of lung tissue homogenates confirmed ample TGF-β neutralization preventing receptor engagement by showing significant inhibition of phosphorylation of Smad 2/3 (60%) (Figs 5B and C), a well-known immediate consequence of engaged TGF-β receptor kinase activity (31). Also, galectin-3, an important regulator of fibrosis and activator of neutrophil mobility (56, 57) was downregulated by 49% (Fig 5B and C). Fibronectin expression, which likely reflects an early inflammatory remodeling phase (58), was downregulated over 80% with 833c&Freso treatment (Figs 5B and C). We did not see any significant blocking of the PI3K/AKT or mTOR signaling pathways or PDGFRβ (Fig S5) (59).

To elucidate key mechanisms and features of the bispecific antibody required to achieve ultra-low dose efficacy, we similarly tested the therapeutic capabilities of 833c, 833cX&Freso and Freso (Fig 6A-C). Neither lung-targeting 833c, which does not carry the therapeutic arm, nor 833cX&Freso, which does not bind to APP2 (Fig 1E), reduced total leucocyte or neutrophil infiltration of BALf at 10 μg/kg doses. Thus, mutating just 2 amino acids in the 833c heavy chain CDR region of the bispecific antibody abolished its APP2 reactivity (Fig 1E), its lung targeting (Figs 2C and D) and its low dose therapeutic potency (Figs 6A-D), while SEC profiles (Fig S1) as well as TGF-β binding (Fig 1F) were maintained. In a separate experiment to rule out possible dual target therapeutic synergy, we also did not detect any infiltration reduction when 833c and fresolimumab were injected together at 10 μg/kg each (Fig S6) whereas 833c&Freso worked well again, indicating that fusion of the two antibodies was essential for therapeutic efficacy. Ultimately antibody binding to APP2 appears to increase potency here not as a direct therapeutic target but rather indirectly as a delivery target mediating caveolae transcytotic pumping into the lung.

**Figure 6.**
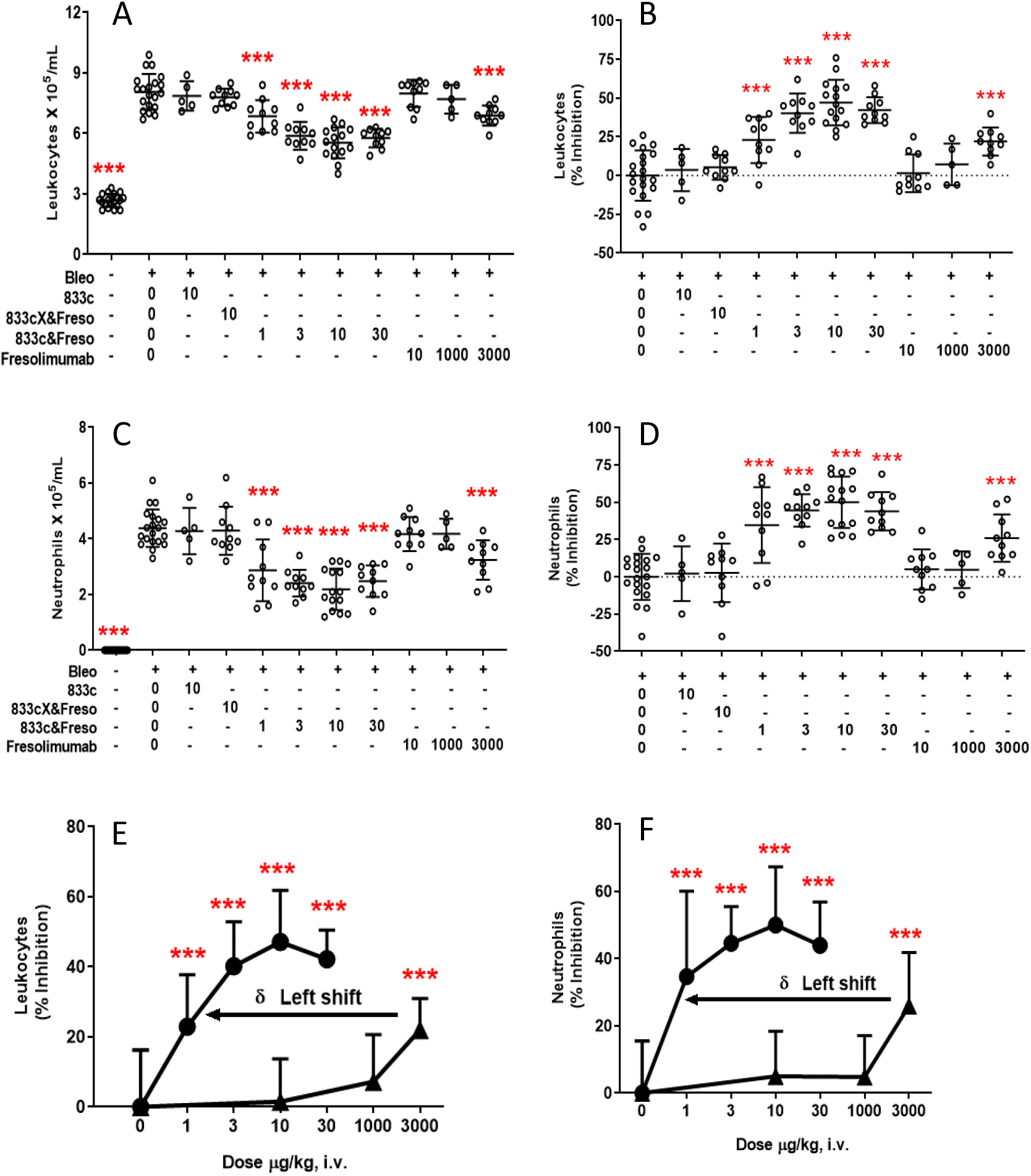
Lung targeting increases therapeutic potency of TGF-β blockade. Two days after bleomycin exposure, rats were treated iv with 833c, 833cX&Freso, 833c&Freso, or fresolimumab at the indicated doses (μg/kg). BALf was collected on day 3. Total leukocytes **(A)** and neutrophils **(C)** per ml of BALf. Percent (%) inhibition of leukocyte **(B**,**E)** and neutrophil **(D**,**F)** accumulation in BALf. **(E**,**F)** 833c&Freso (solid circles), fresolimumab (solid triangles). Graphs and statistical analysis as in Fig 3. Data were pooled from 1 to 4 independent experiments with n=5-20 rats per group.

Fresolimumab also failed at 10 μg/kg dose as well as at 1 mg/kg (Fig 6). It required 3 mg/kg to obtain a statistically significant inhibitory effect (Figs 6A-D). Note the ≥1000-fold increase in potency of 833c&Freso (μg/kg) compared to fresolimumab (mg/kg) (Figs 6E and F). Together these data show that targeted drug delivery via APP2 pumping is required to generate a pronounced therapeutic effect for fresolimumab at μg/kg doses. This precision active lung delivery enables ultra-low dose TGF-β antibody efficacy.

With 70-80% of this novel heterobifunctional therapeutic delivered to the lung in 1 h, caveolae pumping achieves ∼10 times the maximum blood concentration inside the lung (∼10 nM with 10 μg/kg dose). This rapid uptake and concentrating effect, even against an opposing concentration gradient, is consistent with active transport (pumping) into the lung by caveolae. This level inside lung tissue is well above the low nM affinity of the therapeutic arm of the bispecific, fresolimumab. This maximizes immunobinding that neutralizes TGF-β effector functions. Without enough free TGF-β to activate its receptors and downstream signaling, such as Smad phosphorylation (Figs 5B and C), TGF-β’s role in inflammation, edema and biomarkers is inhibited. A 3 μg/kg dose yields near but likely still above Kd concentration, thereby blocking most but not all TGF-β. Our data appear internally self consistent and readily explain why 1 μg/kg but not 3 μg/kg was found statistically less effective than 10 μg/kg.

Fresolimumab on the other hand requires 3 mg/kg iv dose to achieve therapeutic activity (Fig 6). Peak blood levels of ∼300 nM at this dose appear to be needed to drive low nM concentrations required inside the lung to block enough TGF-β to render therapeutic efficacy akin to that observed with the bispecific antibody at 1 μg/kg (Fig 6). Passive transvascular delivery of antibody appears to require at least a 100-fold concentration gradient across the EC barrier to drive enough inside tissue to be effective. In contrast, active transcytosis of APP2 antibody actually creates a gradient in the opposite direction where lung concentrations exceed even peak blood levels (Fig 2) (7). This study exposes just how limiting the EC barrier can be. We discover by overcoming it so readily that the EC barrier in lungs is no longer just theoretical in its impact but rather quite formidable and exigent in limiting Ab uptake and efficacy, even at high doses. With subnM levels in the blood from low μg/kg doses, immunotargeting caveolae pumping achieves therapeutic nM lung concentrations only attainable passively at mg/kg doses. Up to this point the endothelium has been limiting and appeared “unfriendly” to therapy, but now, for the first time, the endothelium through caveolae mediated pumping can be a facilitator promoting therapeutic potency and efficacy.

## Discussion

Like other targets, targeting moieties, and drug delivery pathways and systems discovered with great theoretical promise, our discovery of APP2 and the caveolae pumping system in lung requires stringent, quantitative testing in diseased settings to assess any possible therapeutic benefit. We have begun to report here critical proofs of principle validating the caveolae-mediated drug delivery strategy by retargeting an already exquisitely precise therapeutic drug, the TGFβ blocking antibody fresolimumab. We show that caveolae pumping in the vascular endothelium can still function well in pneumonitis and that caveolae can pump therapeutic antibody into diseased lung to achieve unprecedented precision therapeutic targeting and efficacy, all at strikingly low doses. We have created the first bispecific antibody or other drug that after systemic administration precisely targets inside a single, normally restrictive solid tissue to exert therapeutic efficacy at microdoses. It is engineered to be precisely and actively delivered by caveolae only across the microvascular endothelium in lung tissue. Transendothelial pumping from the blood rapidly concentrates it to levels well above its affinity to help maximize target TGF-β blockade inside the diseased lung tissue and thereby boost its anti-inflammatory potency.

Precision therapeutics are extraordinary in their high affinity and molecular specificity at low concentrations but frequently not in their targeting once injected intravenously. Our therapeutic prototype is the first to be administered systemically (iv bolus) at therapeutic dosages commensurate to the drug’s binding affinity to its target (nM) rather than 2 or more orders of magnitude in excess. High affinity precision antibodies attempting to treat diseases inside a solid tissue usually require μM blood levels attained from standard therapeutic doses of 10 mg/kg or more. The blood levels for small molecule precision therapeutics can easily be 100-fold higher, despite again nM or better affinities. Blood levels near or below 1 nM have never yielded solid tissue targeting and therapeutic efficacy as described herein. That is because passive drug delivery into a tissue with a sufficiently restrictive EC barrier cannot attain interstitial concentrations rivaling or exceeding blood maximums. Passive extravasation is driven not by transcytotic pumping but rather primarily the maximum concentration in the blood which dissipates readily. The caveolae pumping system can continue to transport in a direction opposite to the concentration gradient and thus boosts interstitial concentrations to an order of magnitude above blood levels, all within minutes of iv injection.

Our findings here represent a major advance for therapeutic antibodies, bispecific and in general, towards greater utility and impact in solid tissues with EC barriers limiting passive extravasation. The lung restricts entry and microdose therapeutic efficacy until fresolimumab is re-engineered into a bispecific format enabling retargeting to the caveolae pumping system in lung microvascular endothelium. Once injected iv to circulate in the blood, the two delivery arms of the bispecific antibody have inherent access to its intended EC delivery target in caveolae for active and rapid extravasation to the lung interstitium. Now the two precision therapeutic arms have rapidly attained unprecedented target access so that TGF-β can be comprehensively engaged at high, near saturation concentrations and functionally blocked to abate inflammation at lower doses. The APP2 concentrated in caveolae on the luminal EC surface only in lung microvessels is critical to the precision lung targeting, tissue penetration and concentration process. Without APP2 binding, active drug delivery to lung is lost and so is the dual precision therapeutic efficacy at extraordinarily low μg/kg doses. Standard passive drug delivery ensues and remains limited to requiring >1000-fold higher doses to achieve any efficacy with fresolimumab alone. The precision therapeutic on its own can’t perform optimally without active delivery across the restrictive endothelial barrier.

Getting 70-80% of injected dose into a single solid tissue of the therapeutic antibody or for that matter any drug is remarkable; to do so within 1 hr even more so. Small molecules extravasate passively better than antibodies to bind targets inside restrictive tissues but still have not targeted with such precision, speed, extent and low dosage as 833c&Freso. Usurping physiological pumping mechanisms can concentrate key radioactive small molecules like iodine in thyroid even at low doses but rapid renal filtration excretes the vast majority after iv injection. High affinity ligand-mimicking peptides can bind physiological receptors to accumulate in a few endocrine and tumor tissues but require high blood levels (μM) that dissipate within minutes due to rapid renal elimination (60, 61).Moreover, unlike many past studies reporting preclinical testing of mono-, bi- and tri-specific antibodies including some reaching clinical trials (62, 63), all results here occur under native in vivo conditions with only endogenous target expression levels and are devoid of any target and/or species-specific manipulations rendering favor to the desired delivery and therapeutic outcome. Active transvascular delivery can indeed enable precision targeting well beyond passive delivery.

Here we show active delivery by targeting caveolae-mediated transcytosis in lung improves precision targeting by >100-fold which leads to even greater enhancement in precision therapeutic potency. Targeting and therapeutic goals near theoretical expectations can be achieved with a bispecific antibody at nM concentrations - all to the benefit of avoiding toxicities. Furthermore, the rapid blood clearance (∼80% in 1hr) not from metabolism or renal excretion but rather robust uptake in a single desired target tissue minimizes exposure and toxicities elsewhere in the body.

Although representing a major step forward towards achieving “magic bullet” utility, this study uses only a single therapeutic construct in one disease model. This study and the caveolae-mediated drug delivery paradigm in general have many unaddressed issues and possible limitations. Establishing broader utility in other lung diseases awaits further study. Some pathologies may more directly affect endothelial cell function to possibly disable caveolae pumping. Whether caveolae targeting works in other organs, normal and diseased, requires more target discovery. Also, the bispecific format used here is limited; it is only one of many and provides only two blocking opportunities per molecule. Loading antibodies with small inhibitors either directly via chemical conjugation or indirectly for instance by attaching drug nanocarriers, may provide more impact. Utilizing antibody derivatives and other targeting agents to study the effects of size, affinity, avidity, drug loading and nanoparticles on caveolae pumping and precision tissue targeting may even provide further insights and improvements beyond an already prodigious start. So far, we do not detect any evidence of interference by target presence in circulating blood, for instance from APP2 shedding and/or TGF-β secretion. In human serum, TGF-β is sequestered and functionally inactive in the blood protein alpha-2 macroglobulin (64). However, this could become potentially limiting in other pathologies.

This new approach provides the first benefits bestowed through a new active drug delivery paradigm; a means to penetrate the EC barrier that has plagued delivery into many normal and diseased tissues/organs; a means to more readily and fully engage therapeutic targets usually hidden inside so many organs. We have created a new class of “dual precision” medicine by coupling two specific antibodies, one providing unprecedented lung delivery, the other exclusive therapeutic target neutralization. The result is a highly specific molecular therapeutic agent that elevates the *in vivo* pharmacological potency by 3 orders of magnitude directly by solving the delivery problem. Lung targeting via caveolae appears paramount to achieve the marked therapeutic effect of 833c&Freso at μg/kg doses.

We ultimately demonstrate the therapeutic value of accurate single tissue drug delivery and penetration. Overcoming the EC barrier through caveolae pumping may be key to maximizing therapeutic impact and eliminating ambiguity as to cause when a drug fails. If delivery is assured, then the drug’s pharmacological activity can be maximized to yield efficacy commensurate with the target effector’s role in the disease. Dual delivery and therapeutic precision targeting encourages a paradigm shift from passive to active drug delivery; accordingly therapeutic drug doses may drop from mg/kg to μg/kg. Many drugs may benefit from similar targeted caveolae pumping, especially drugs showing promising activity yet limited by high dose toxicities, sometimes even leading to failed clinical trials. This novel delivery strategy could expand our applicable drug repertoire and ultimately revolutionize treatment options, if successful in clinical trials.

## Author contributions

Conceptualization: AHK, JES

Methodology: AHK, KK, TB, BC, CY, JR, MDL, JK, SN, BO, AC, JES

Funding acquisition: ZB, JES

Writing – original draft: AHK, KK, JES Writing – review & editing: ZB, JES

## Supplementary Figures

**Figure S1.**
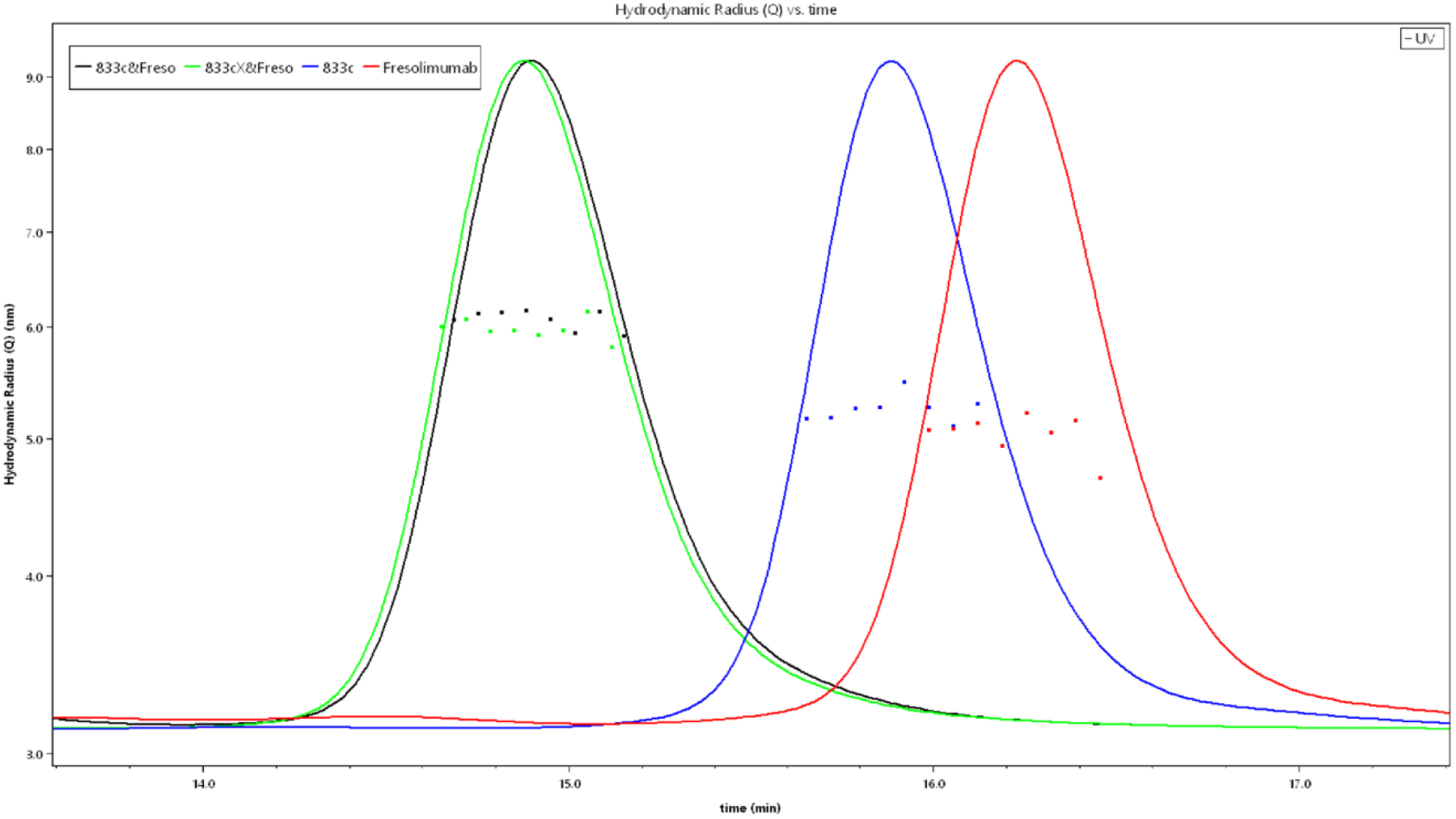
Size-exclusion chromatogram with multi angle light scattering detection of 833c, fresolimumab, 833c&Freso and 833cX&Freso. Average hydrodynamic radius (nm): 833c=5.2, fresolimumab=5.0, 833c&Freso=6.1, and 833Xc&Freso=6.1.

**Figure S2.**
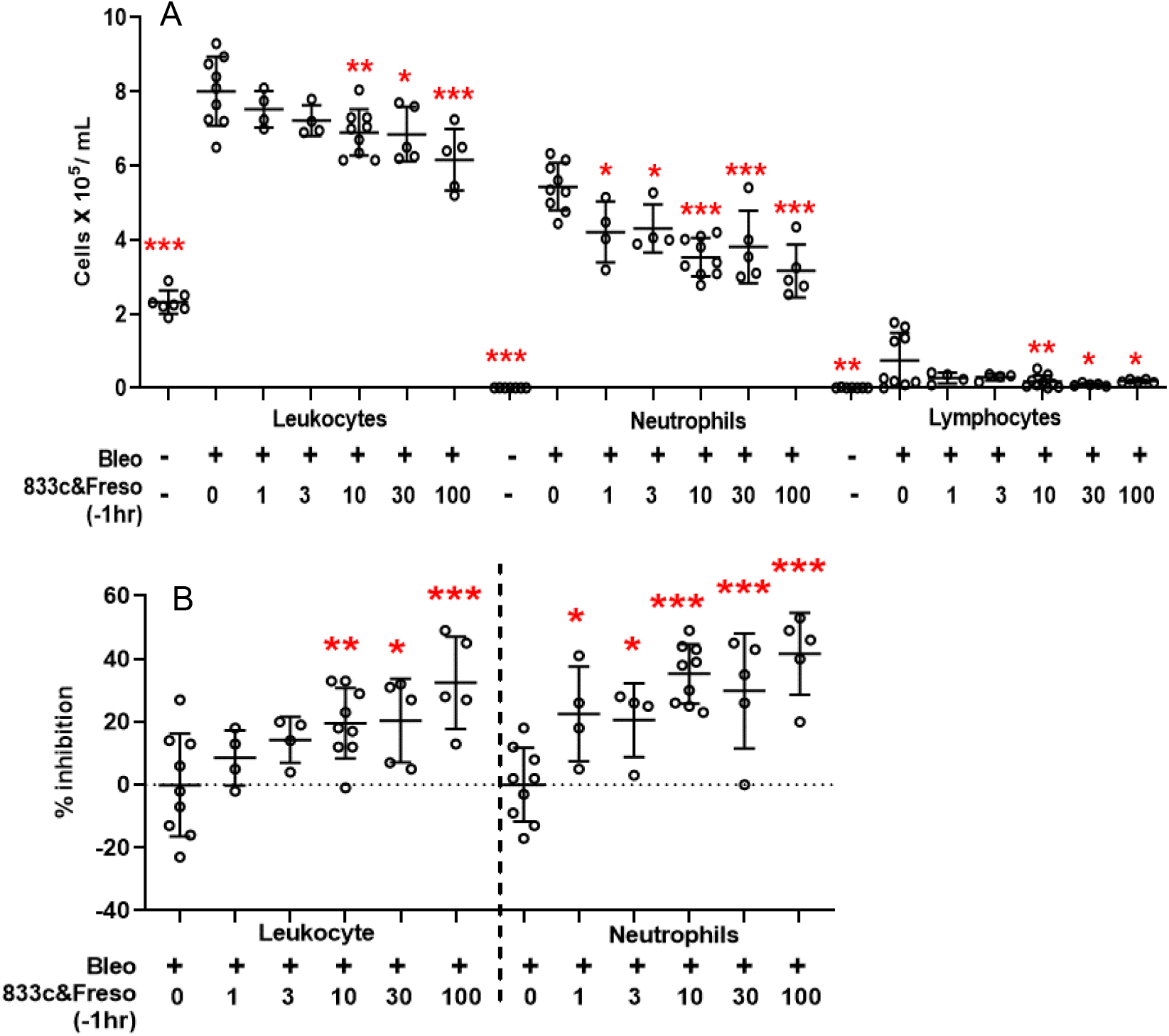
Prophylactic effect of 833&Freso on inflammatory cells in bronchoalveolar fluid (BALf). Rats were treated with 833c&Freso at the indicated doses (μg/kg) by iv injection one hour before (−1 h) administering bleomycin (bleo) it (see Methods). BALf was collected on day 3. **(A)** Total and differential leukocyte concentrations in BALf. **(B)** Percent (%) inhibition of total leukocyte and neutrophil infiltration. Data were pooled from 2 independent experiments with 833c&Freso 10 μg/kg repeated in both experiments (n=4-9 total rats per group). Scatter graphs of each data point are shown with indicated means ± SD. *P< 0.03, **P< 0.01, and *** P< 0.001 vs. bleo only, by using one-way ANOVA followed by Dunnett’s test.

**Figure S3.**
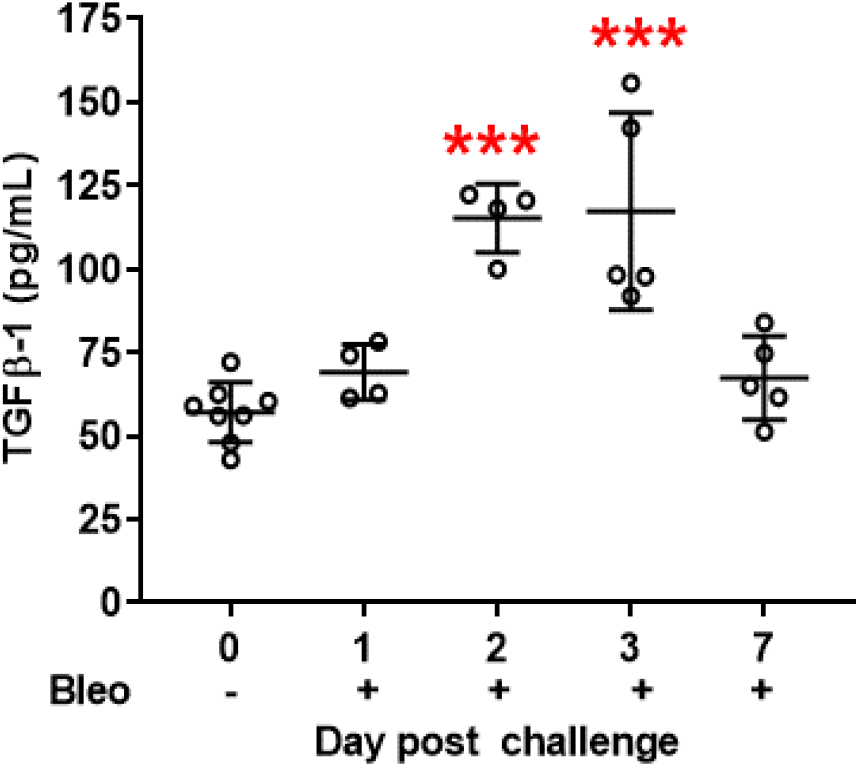
TGF-β expression in bleomycin challenged rats. Rats received a single it instillation of bleomycin as described in Methods. BALf was collected and TGF-β1 concentration determined by ELISA. ***P<0.001 vs day 0, by using one way ANOVA followed by Dunnett’s test.

**Figure S4.**
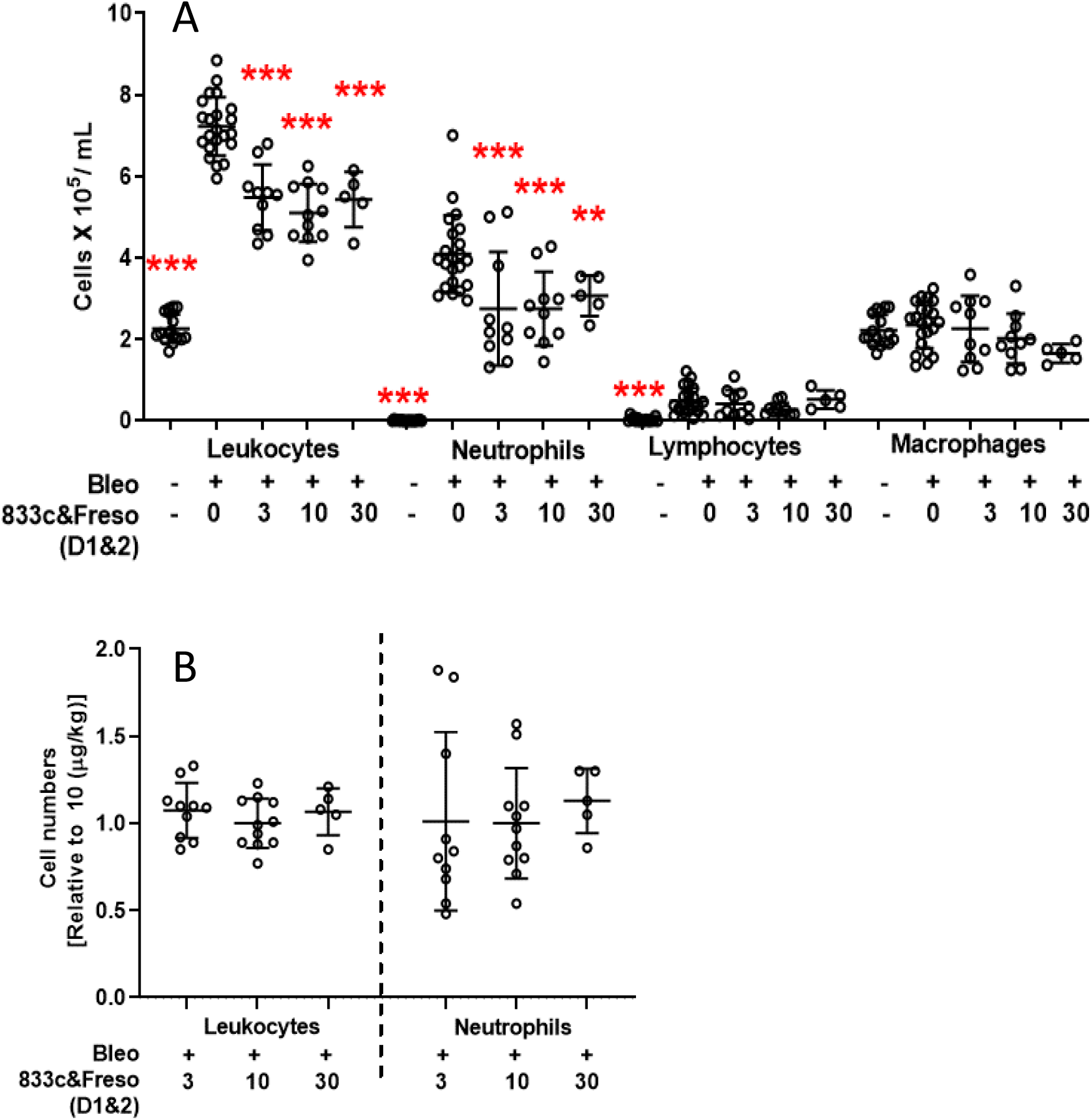
Dose optimization of 833c&Freso. Rats were injected with 833c&Freso at the indicated doses (μg/kg) on day 1 and day 2 (D1&2) after bleomycin (bleo) exposure. BALf was collected on day 3. **(A)** Total and differential leukocyte profile in BALf. **(B)** Relative cell numbers normalized to 10 μg/kg. Data pooled from 1-4 independent experiments with n=5-22 total rats per group. Scatter graphs of each data point with indicated means ± SD. *P< 0.03, **P< 0.01, and *** P< 0.001 vs. bleo only, by using one-way ANOVA followed by Dunnett’s test.

**Figure S5:**
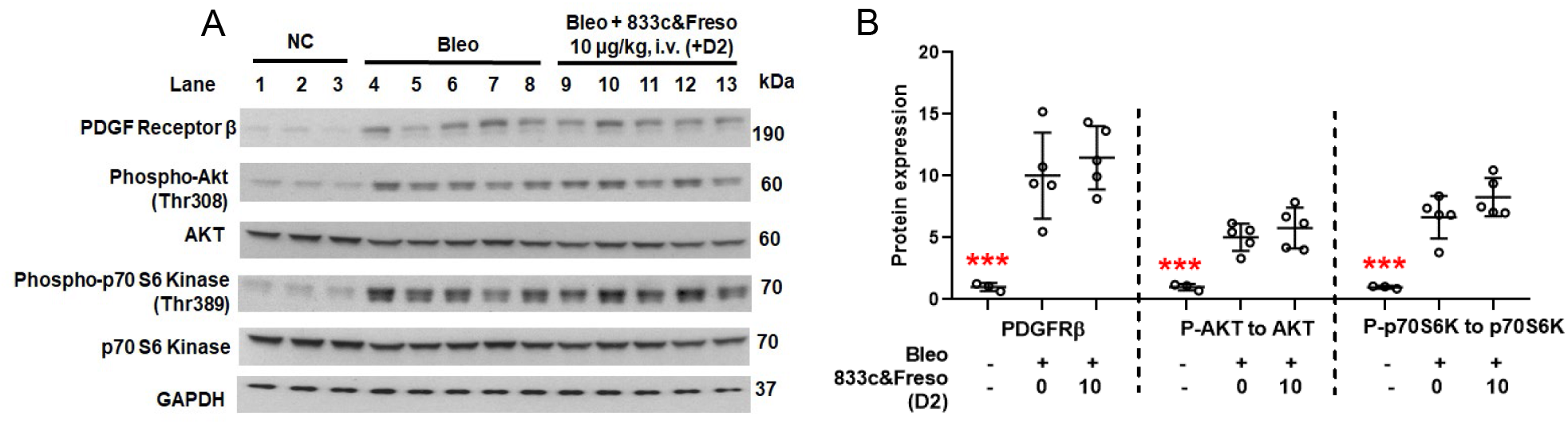
833c&Freso does not affect PDGF/AKT/mTOR axis signaling in the lung. Rats were treated with 10 μg/kg 833c&Freso iv on day 2 post-bleomycin. Lung tissue was harvested on day 3. **(A)** Lung tissue homogenates were subjected to western blot analysis using antibodies to indicated proteins. **(B)** Relative expression of PDGFRβ, P-AKT to AKT and P-p70S6K to p70S6K derived from western blot image analysis. Scatter graphs of each data point with indicated means ± SD. N=3-5 rats per group. *** P< 0.001 vs. bleo only, by using one-way ANOVA followed by Dunnett’s test.

**Figure S6.**
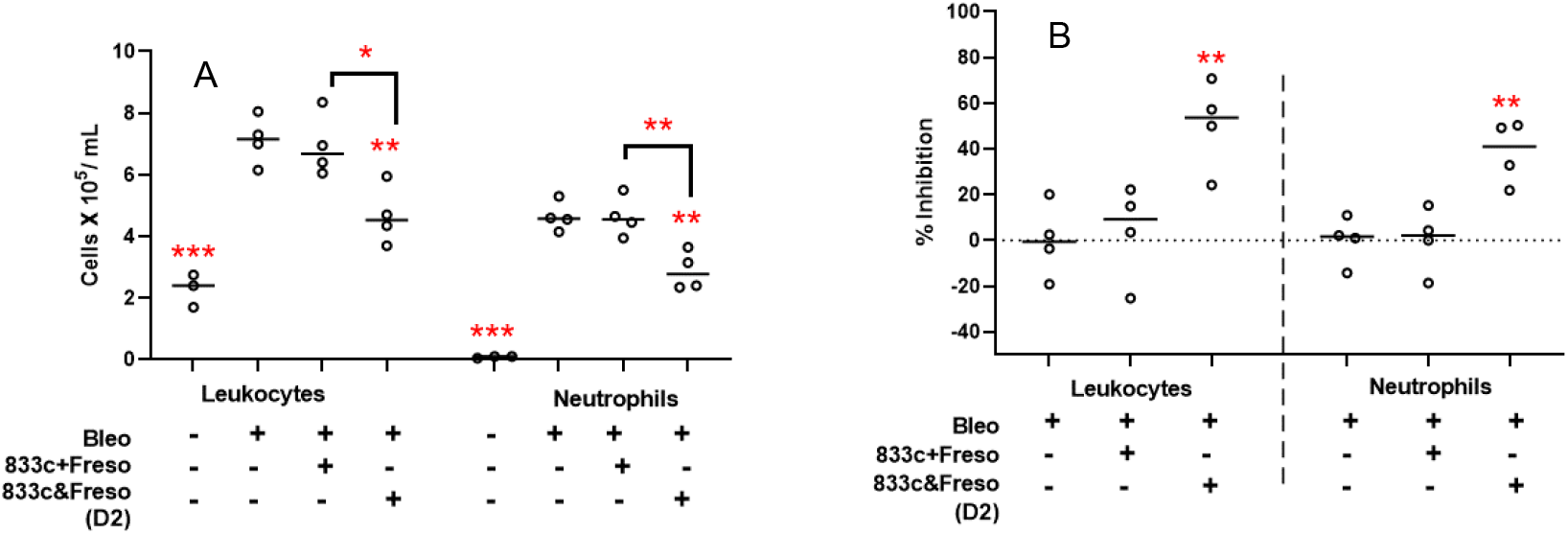
833c plus fresolimumab does not affect leukocyte and neutrophil infiltration. Rats were injected with either10 μg/kg of 833c and fresolimumab or 10μg 833c&Freso on day 2 after bleomycin (bleo) exposure. BALf was collected on day 3. **(A)** Total leukocyte and neutrophils in BALf. **(B)** Percentage (%) inhibition of leukocyte and neutrophil infiltration. N=4 rats per group. Scatter graphs of each data point with indicated means ± SD. *P< 0.03, **P< 0.01, and *** P< 0.001 vs. bleo only, by using one-way ANOVA followed by Dunnett’s test.

